# Consequences of daily rhythm disruption on host-parasite malaria infection dynamics

**DOI:** 10.1101/2023.08.24.554632

**Authors:** Jacob G. Holland, Kimberley F. Prior, Aidan J. O’Donnell, Sarah E. Reece

**Affiliations:** Institute of Ecology and Evolution, University of Edinburgh, Edinburgh, UK; Institute of Immunology and Infection Research, University of Edinburgh, Edinburgh, UK

**Keywords:** Circadian rhythm, Plasmodium, intra-erythrocytic development cycle, fitness, virulence, transmission

## Abstract

Undertaking certain activities at the time of day that maximises fitness is assumed to explain the evolution of circadian clocks. Organisms often use daily environmental cues such as light and food availability to set the timing of their clocks. These cues may be the environmental rhythms that ultimately determine fitness, act as proxies for the timing of less tractable ultimate drivers, or are used simply to maintain internal synchrony. While many pathogens/parasites undertake rhythmic activities, both the proximate and ultimate drivers of their rhythms are poorly understood. Explaining the roles of rhythms in infections offers avenues for novel interventions to interfere with parasite fitness and reduce the severity and spread of disease. Here, we perturb several rhythms in the hosts of malaria parasites to investigate why parasites align their rhythmic replication to the host’s feeding-fasting rhythm. We manipulated host rhythms governed by light, food, or both, and assessed the fitness implications for parasites, and the consequences for hosts, to test which host rhythms represent ultimate drivers of the parasite’s rhythm. We found that alignment with the host’s light-driven rhythms did not affect parasite fitness metrics. In contrast, aligning with the timing of feeding-fasting rhythms may be beneficial for the parasite, but only when the host possess a functional canonical circadian clock. Because parasites in clock-disrupted hosts align with the host’s feeding-fasting rhythms and yet derive no apparent benefit, our results suggest cue(s) from host food act as a proxy rather than being a key selective driver of the parasite’s rhythm. Alternatively, parasite rhythmicity may only be beneficial because it promotes synchrony between parasite cells and/or allows parasites to align to the biting rhythms of vectors. Our results also suggest that interventions can disrupt parasite rhythms by targeting the proxies or the selective factors driving them without impacting host health.

## 1. INTRODUCTION

Biological clocks allow organisms to adaptively respond to predictable rhythmic changes in their environments (Yerushalmi & Green, 2009). The environment provides many potential rhythmic cues across the day, such as light, temperature, humidity, tides, and the rhythms of other organisms (Helm et al., 2017) which can act as inputs to clocks (‘Zeitgebers’). These time cues may themselves directly impact fitness or else provide information about correlated cyclical opportunities and risks that impact fitness, or may simply facilitate beneficial synchronisation of processes within the organism regardless of the consequences of environmental alignment (‘intrinsic advantage’, Krittika & Yadav, 2020). Thus, an organism can schedule a rhythmic activity to align with an environmental rhythm because the environmental oscillation directly impacts on the organisms’ fitness, provides convenient timing information, or both. Understanding how environmental rhythms shape activities therefore requires disentangling their role as time cues from their roles as selective forces that drive the evolution and maintenance of rhythms (Hut & Beersma, 2011). Assuming an organism’s rhythms are adaptive (i.e., enhance fitness), this can be achieved by perturbing the alignment of an organism’s rhythms to various environmental rhythms to ascertain which perturbations have detrimental fitness implications. However, this is challenging because the links between environmental inputs, endogenous clocks, clock-controlled outputs, and fitness returns are often unknown. Moreover, organisms often utilise multiple Zeitgebers (Harder & Oster, 2020), many of which interact, producing an array of rhythmic outputs which themselves may be regulated by multiple inputs, increasing the challenges of disentangling the fitness impacts of following particular environmental rhythms.

The use of cues to determine biological processes has long been studied in the context of phenotypic plasticity (reviewed in Schneider, 2022; Snell-Rood & Ehlman, 2021), leading to the distinction between cues which are the ‘selective drivers’ of evolution, i.e., environmental changes with direct fitness impacts, and cues which are ‘proxies’, i.e., correlated with the selective driver but not of fitness significance *per se*. Importantly, proxies act as less reliable cues because they may not always perfectly reflect the state of selective drivers, but may be more convenient to measure; for example, various prey animals use light or temperature as a correlate for predation risk (Miehls, McAdam, Bourdeau, & Peacor, 2013; Orrock, Danielson, & Brinkerhoff, 2004; Suppa et al., 2021). Likewise, in a circadian context, the cues causing (‘effecting’) rhythmicity may be either the ultimate factors driving the evolution of rhythmicity or just correlative proxies. For example, the diel light cycle (day and night), which acts as a primary Zeitgeber for the circadian clocks of many organisms, may directly impact fitness (e.g. via photosynthetic potential, visual acuity or desiccation risk), but also correlates with changes in the activity of other organisms, generating the potential for a wide range of social or exploitative interactions (positive and negative). Time cues can also help to temporally separate mutually interfering processes (e.g. Chen, Odstrcil, Tu, & McKnight, 2007) or to maintain homeostasis by synchronising different rhythmic processes across levels of biological organisation (Vaze & Sharma, 2013). In this context, the time of day processes are undertaken does not directly impact fitness but following a time cue is a convenient way to ensure separation or synchronisation of processes.

Ascertaining to what extent the proximate cues that effect rhythmic activities are also ultimate drivers offers a novel approach to managing and manipulating interactions between organisms. For example, rhythms in the activities of parasites (including pathogens) underpin their virulence and transmission, whereas rhythms in host immune responses and feeding patterns can determine the severity and outcome of infections. Understanding the links between responses to cues and fitness consequences in rhythmic parasite activities, and in host defences, are fundamental steps towards reducing the benefits parasites garner from their rhythms and/or harnessing host rhythms to control infections (Hunter, Butler, & Gibbs, 2022; Westwood et al., 2019). For example, *Trypanosoma brucei*, which causes sleeping sickness, uses host body temperature to entrain its circadian clock which, in turn, coordinates the expression of its metabolism-related genes (Rijo-Ferreira, Pinto-Neves, Barbosa-Morais, Takahashi, & Figueiredo, 2017). Intuition suggests this allows the parasite to coordinate its own feeding with that of its host, but whether the host’s feeding-fasting rhythms are the ultimate driver remains untested. Rhythmic replication by malaria parasites also aligns with the host’s feeding-fasting rhythms. Malaria (*Plasmodium*) parasites are famously rhythmic; completing cycles of replication with the host’s red blood cells (RBCs) at 24, 48, or 72 hours, depending on the species (Dos Santos, Pereira, & Garcia, 2021; Garcia, Markus, & Madeira, 2001; Mideo, Reece, Smith, & Metcalf, 2013). Each cycle – termed the intraerythrocytic developmental cycle (IDC) – culminates in synchronous bursting to release progeny that initiate the subsequent round of RBC invasions, causing the periodic fever that characterises malaria infection (Gazzinelli, Kalantari, Fitzgerald, & Golenbock, 2014). In the rodent malaria model *P. chabaudi*, the timing of transitions between IDC stages aligns to host rhythms associated with feeding-fasting, even when the host’s light-dark cycle is in an opposing phase or when the host’s canonical circadian clock machinery (transcription-translation feedback loop; TTFL) is disrupted (Hirako et al., 2018; O’Donnell, Prior, & Reece, 2020; O’Donnell, Greischar, & Reece, 2022; Prior et al., 2018). IDC completion occurs towards the end of the feeding window, which is night-time for nocturnally active rodent hosts. The IDC rhythm is at least in part under the control of parasite genes (Prior et al., 2020; Rijo-Ferreira et al., 2020; Subudhi et al., 2020), and if its timing is perturbed, the IDC speeds up by 2-3 hours per cycle until realigned to host rhythms (O’Donnell et al., 2022). The IDC rhythm is important for parasite fitness; it maximises within-host replication, results in transmission stages (gametocytes) being at their most infectious during the night time when mosquito vectors forage for blood, and confers tolerance to antimalarial drugs (O’Donnell, Greischar, & Reece, 2022; O’Donnell, Schneider, McWatters, & Reece, 2011; Owolabi, Reece, & Schneider, 2021; Pigeault, Caudron, Nicot, Rivero, & Gandon, 2018; Schneider et al., 2018).

It is not known whether aligning with rhythms effected by the host’s feeding-fasting schedule provides a direct fitness benefit to parasites and/or simply provides a convenient cue (or series of cues) for the timing of other host/vector rhythms that impact parasite fitness, or whether there are intrinsic benefits of rhythmic replication. For example, a variety of rhythmic host processes are entrained by light, as well as feeding, and these can be mediated in different tissues by the TTFL clock machinery (Astiz, Heyde, & Oster, 2019; S. Zhang et al., 2020). Rhythms in immune responses are often correlated with the timing of feeding-fasting, although immune defences are unlikely to impose the IDC rhythm by killing mis-timed parasites (Cabral, Tekade, Stegeman, Olivier, & Cermakian, 2022; Hunter et al., 2022; Prior et al., 2020). Aligning with feeding-fasting rhythms could directly impact on parasite replication because the host’s digestion of food regulates when parasites have access to essential nutrients they cannot scavenge from haemoglobin. These resources include vitamins B1 and B5, folate, purines, and the amino acid isoleucine (which is absent from human haemoglobin and uniquely rare in murine haemoglobin), that are required by later IDC stages for biogenesis (Skene et al., 2018). Indeed, recent work reveals that as well as being an essential rhythmically available resource, blood isoleucine concentration also fulfils criteria of a time cue used by *P. chabaudi* to set its IDC schedule (Prior et al., 2021). This suggests that isoleucine could be both a proximate cue and an ultimate driver; parasites respond to isoleucine rhythms because isoleucine availability regulates replication, and the isoleucine rhythm is in phase with other nutrients that are most easily acquired from the host’s food. Whether it is also beneficial to align gametocyte development with rhythmic nutrients is not known, and feeding-fasting rhythms may alternatively, or additionally, be a proxy for vector activity rhythms (i.e. because both are correlated with the day-night cycle). According to the intrinsic benefits hypothesis, synchrony of the IDC might be beneficial *per se*, irrespective of external rhythms, and parasites might use time cues to simply synchronise development or coordinate life history decisions (e.g. cell-cell communication involved in reproductive investment decisions (Schneider & Reece, 2021). However, theory also predicts that if too tightly synchronised, parasites inadvertently compete with each other for resources (Greischar, Read, & Bjornstad, 2014), which is supported by the observation that *P. chabaudi* performs better when desynchronised (Owolabi et al., 2021). Thus, both the timing and synchrony of the IDC could independently impact fitness.

Here, we investigate to what extent the host’s light-driven and feeding-fasting rhythms are ultimate drivers of the IDC rhythm by comparing parasite performance in hosts in which: 1) the timing of feeding-fasting rhythms is matched or mismatched (12h out of phase) to light-dark rhythms; 2) feeding is restricted to 12h windows or available throughout the day; and 3) TTFL clocks are disrupted and feeding rhythms are either naturally attenuated or experimentally imposed. We also investigated whether the severity of disease symptoms experienced by hosts depends on how their parasites perform, their own rhythms, and their access to food. We found that aligning with the host’s light-dark rhythms is not an ultimate driver of parasite rhythms, and that any fitness benefits of aligning with the host’s feeding-fasting rhythms requires feeding rhythms to be naturally spread-out and accompanied by a functional TTFL clock. Determining the fitness consequences of rhythms for parasites and hosts is timely and important given that plasticity in the IDC schedule helps parasites tolerate antimalarial drugs (Teuscher et al., 2010), and that the temporal selective landscape of malaria parasites is changing because malaria-vectoring mosquitoes are evading bed nets by altering the time of day they forage for blood (Thomsen et al., 2017). Furthermore, beyond malaria, understanding why the timing and synchrony of parasite replication are ultimately connected to the daily rhythms of hosts may make drug treatment more effective and less toxic to patients.

## 2. METHODS

### 2.1 Hosts and parasites

We used both C57BL/6 J (WT) wildtype and *per1/2*-null clock-disrupted mice backcrossed onto a C57BL/6 J background for over 10 generations (O’Donnell et al., 2020). All experimental mice were females, approximately 11 weeks-old at the start of the experiment and had been group housed at ∼ 20 oC, 60% RH, with a 12:12 Light: Dark regime (lights on 0800-2000; all times in UCT+1), *ad libitum* access to food (RM3 pellets, 801700, SDS, UK) and unrestricted access to drinking water supplemented with 0.05 % para-aminobenzoic acid (Jacobs, 1964). Two weeks before infection, we singly housed mice and randomly allocated them to treatment groups (n =5 per group) to begin their experimental photoschedules and feeding treatments, which we maintained for the duration of the experiment. All wildtype mice remained in LD 12:12, in which they exhibit nocturnal activity and foraging via the mammalian circadian system; the TTFL clock oscillates in cells throughout the body and keeps time via entrainment to light (Finger & Kramer, 2021; Partch, Green, & Takahashi, 2014). In contrast, *per1/2*-null mice were transferred to constant darkness in which they are behaviourally arrhythmic because null versions of the *Per1* and *Per2* clock genes disrupt the canonical TTFL machinery, and no light-dark cues are available to invoke direct (‘masking’) responses (Bae et al., 2001; Maywood, Chesham, Smyllie, & Hastings, 2014; O’Donnell, Prior, & Reece, 2020). All mice received an intravenous infection of 10^5^ red blood cells infected with *P. chabaudi* (genotype DK) at the ring stage, an early stage in the IDC. DK parasites cause relatively mild infections, which minimises off-target effects of sickness on host rhythms and parasite performance (Prior et al., 2019).

### 2.2 Experimental design

We used five treatment groups to compare parasite and host performance metrics within pairs of groups that enabled three questions to be addressed (Figure 1). Our approach aimed to decouple the time cues available to parasites from different host rhythms in manners that minimise confounding impacts of forcing parasites to alter IDC schedule which would occur if treatments involved misaligning parasites to feeding-fasting rhythms. The treatments were: (i) “WT-AL” (Wild Type - *ad libitum*), wild type hosts in LD with food constantly available. The feeding-fasting and light-entrained rhythms of these mice are aligned because they follow their natural patterns of nocturnal behaviour and undertake the bulk of their foraging in the dark; (ii) “WT-DF” (Wild Type – Dark Fed), wild type hosts in LD with time-restricted feeding in which food was only available during the dark period of each circadian cycle. The feeding-fasting and light-entrained TTFL rhythms of these mice are aligned and they differ from the WT-AL group because they cannot eat between dawn and dusk. Mice experiencing TRF consume approximately the same amount of food per day (after an initial adjustment period) as *ad libitum* fed mice, even when the window available for feeding is restricted to a few hours (e.g. Froy, Chapnik, & Miskin, 2006; Hatori et al., 2012), including when infected with malaria (O’Donnell et al., 2020). (iii) “WT-LF” (Wild Type – Light Fed), wild type hosts in LD with time-restricted feeding in which food was only available during the light period of each circadian cycle. By inverting the timing of food availability relative to the light-dark schedule, the parasite’s IDC continues to be aligned to the host’s feeding-fasting rhythms but becomes misaligned to the host’s light-entrained rhythms. (iv) “*per1/2*-RF” (*per1/2*-null - Restricted Fed), *per1/2*-null hosts in DD with time-restricted feeding in which food was only available during a 12-hour window each day (2000-0800). These hosts exhibit experimentally imposed rhythmicity in some processes related to the digestion of food, metabolism, and fasting, but with no influence of TTFL-driven clocks. (v) “*per1/2*-AL” (*per1/2*-null *ad libitum*), *per1/2*-null hosts in DD with food constantly available. These mice are essentially arrhythmic, exhibiting short and frequent bouts lasting minutes of feeding (O’Donnell et al., 2020), and thus offer no rhythmic time cues to parasites. To ensure that all infections were initiated with parasites aligned with the feeding-fasting schedule of their recipient host, we harvested parasites from donor hosts housed in two different LD 12:12 photoschedules. Specifically, parasites were harvested from the end of the respective donor dark periods at 0830 to infect the WT-AL, WT-DF, *per1/2*-RF and *per1/2*-AL treatments, and at 2030 on the same day to infect the WT-LF treatment. We checked food twice daily to ensure a constant supply to *ad libitum* fed mice (WT-AL, *per1/2*-AL), and swept cages for stray pellets when food was removed from TRF mice (*per1/2*-RF, WT-DF, WT-LF).

**FIGURE 1.**
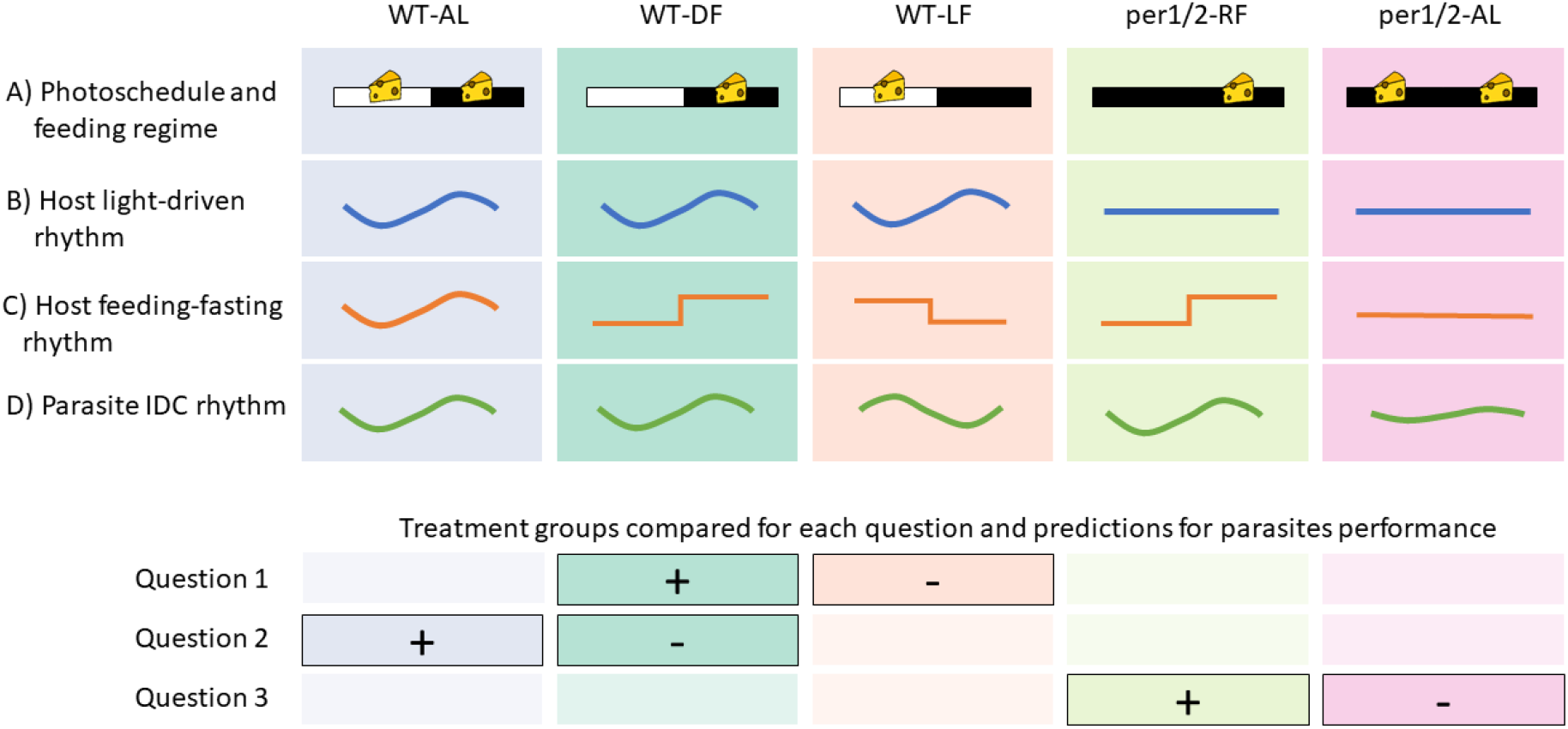
Characteristics of treatment groups and rationale for study design. The upper four rows show: (A) Photoschedule; either light-dark cycles (LD12:12, white and black bars) or constant darkness (DD, black bars). Feeding regime; time restricted feeding limited to 12 hours per day (RF, 1 cheese) or *ad libitum* (AL, 2 cheeses). How the treatment groups differ in terms of the typical pattern and timing of: (B) light-driven TTFL rhythms governed by the SCN, illustrated by locomotor activity, (C) feeding-fasting and downstream peripheral rhythms, illustrated by feeding, and (D) the parasite’s IDC schedule, illustrated by the timing of bursting to release progeny. The IDC rhythm can be decoupled from rhythms entrained by the host’s light-dark cycle, but consistently reschedules to the host’s feeding-fasting rhythm, precluding direct assessment of the fitness impacts of feeding-fasting associated rhythms. The lower four rows show the pairwise comparisons between treatments (solid boxes) used to test each of the following key questions, where -/+ denote the treatments in which parasites are expected to perform worse/better within each pair, respectively: **(Q1) Do parasites derive ultimate fitness benefits from aligning to host rhythms entrained by the light-dark cycle?** If feeding-fasting cues are used as a proxy for light-entrained rhythms that impact on parasite fitness, parasites will perform worse when aligned to feeding-fasting rhythms that are decoupled from light-entrained rhythms (WT-LF). **(Q2) Do parasites derive greater fitness benefits when rhythmic hosts have typically rhythmic feeding-fasting?** *Ad lib* fed hosts spread their food intake around a peak in the dark phase (O’Donnell et al., 2020), thus, feeding cues may peak at similar times in WT-AL and WT-DF hosts but *ad lib* hosts take in food over a longer window that includes dusk and dawn. If the IDC rhythm represents a balance between the benefits of timing to align with nutrient availability versus the costs of extreme synchrony causing competition, this constraint will be ameliorated in WT-AL hosts who can spread their feeding out and so we predict that parasites will perform better in WT-AL hosts. **(Q3) Do parasites benefit from specifically aligning to feeding-fasting rhythms in TTFL-disrupted hosts with no other discernible rhythms?** Parasites align to feeding-fasting rhythms even in clock disrupted hosts (via TRF). If parasites benefit from non-TTFL mediated aspects of rhythmic host feeding (Greenwell et al., 2019; O’Donnell et al., 2020) or from intrinsic benefits of synchrony, and these benefits outweigh potential costs, parasites will also perform better in *per1/2*-RF than *per1/2*-AL hosts. Finally, we also predicted that in groups in which parasites performed better, hosts will experience more severe infection symptoms, but that hosts with constant access to food (WT-AL and *per1/2*-AL) can cope better with infection.

### 2.3 Sampling and data collection

Previous studies have attempted to assess fitness impacts from only a few IDCs at the start of infections (e.g. O’Donnell, Mideo, & Reece, 2013), but the selective advantage of the IDC may vary throughout infection (Prior et al., 2020). To overcome this limitation, we monitored infections throughout the acute phase which includes recovery from the peak of infections and captures the bulk of gametocyte production to assess transmission potential. Specifically, we assessed parasite performance in terms of overall parasite and gametocyte dynamics, and infection severity in terms of anaemia and weight loss. We sampled mice daily from day 3 to day 17 post infection (PI) at 0830 for the *per1/2*-RF, *per1/2*-AL, WT-AL and WT-DF treatments, and at 2030 for the WT-LF treatment, to ensure the age of infection (in hours) was consistent across treatments. Four mice were euthanised due to reaching the humane endpoints of infection in the following treatments (at days PI): WT-DF (10), *per1/2*-RF (8), *per1/2*-AL (9, 9). At each sampling point, we weighed the mice and collected blood samples (2µl for RBC density, 5µl for total parasite density, and 10µl for gametocyte density).

We measured RBC density using a particle counter (Beckman Coulter Z2). For total parasite density, we mixed 5 µl blood samples with 150 µl citrate saline upon collection and (after centrifuging and discarding the plasma supernatant) extracted DNA for qPCR. For gametocyte density, we mixed 10 ul blood samples with 20 µl RNAlater® upon collection, and extracted RNA for RT-qPCR. We followed extraction and qPCR protocols targeting the CG2 gene (PCHAS_0620900) as detailed elsewhere (Owolabi et al., 2021; Petra Schneider et al., 2015). Notably, since the CG2 gene is expressed only in gametocytes (Wargo, De Roode, Huijben, Drew, & Read, 2007), CG2 cDNA quantifies the number of gametocytes, whereas CG2 DNA quantifies the total number of asexually replicating stages and gametocytes.

### 2.4 Data analysis

We quantified metrics for parasite fitness using parasite density as a measure of in-host survival, and gametocyte density as a measure of transmission potential. For each of these density metrics we analysed: i) the dynamics of log-transformed density throughout the infection, using day PI as a factor (since density is non-linear) and random intercepts for each mouse ID; ii) peak density, defined as the highest log-transformed density observed for each infection (or each infection wave for gametocytes); and iii) overall density, defined as the cumulative number of parasites observed throughout infection and calculated only from mice that survived the entire experiment. For parasite density, two samples were defective and excluded. For gametocytes, the peak density was analysed separately for the ‘early’ and ‘late’ waves of each infection (before or after day 10PI), because *P. chabaudi* exhibits two peaks of gametocytes. We excluded *per1/2*-RF infections from all gametocyte analyses due to loss of the RNA samples for this group, meaning it was not possible to test Q3 in relation to transmission potential. We quantified weight loss and anaemia as the difference between weights and RBC densities on day 3PI and at their respective troughs.

All analyses were conducted using R version 4.0.0 or later (R core development team 2020). We took a two-stage approach to the analysis of each metric. First, we produced a separate linear model or linear mixed model (using the lme4 package v 1.1.32; Bates, Machler, Bolker, & Walker, 2015) set for each metric as a response variable and checked assumptions using the DHARMa package (v 0.3.3.0; Hartig, 2020). We then compared whether each of these models were more parsimonious (lower AICc) than the respective null models (i.e. with no treatment term). Second, for those models where metrics varied detectably between treatments (without interaction), we conducted pairwise comparisons corresponding to our three main questions (Figure 1). We configured the models with the reference group (model intercept) as the WT-DF treatment for contrasts with WT-LF (Q1) and WT-AL (Q2), and with the reference group as the *per*1/2-AL treatment for contrast with *per*1/2-RF (Q3), and examined individual contrasts between these levels. For linear mixed models, we estimated p-values via Satterthwaite’s degrees of freedom method using the ‘lmerTest’ package (v 3.1.3; Kuznetsova, Brockhoff, & Christensen, 2017). We estimated effect sizes and confidence intervals for figures using nonparametric bootstrap resampling via the dabestr package (v 0.3.0; Ho, Tumkaya, Aryal, Choi, & Claridge-Chang, 2019).

## 3. RESULTS

### 3.1 Parasite density

For the dynamics of parasite density, the most parsimonious model only included treatment and day PI (model weight = 0.80, delta AICc of all other models > 3.08, see Table S1), implying that the dynamics of all groups followed a similar trajectory over time but varied in magnitude (Figure 2A, B). Pairwise comparisons within this model only revealed a difference in the comparison for Q1, but in the opposite direction to the prediction. Specifically, relative to WT-DF infections, the density was 43% higher in WT-LF infections (Q1; t = 2.4, df = 341, p = 0.016, coefficient of log density = 0.36, 95% CI 0.21-0.51), but did not differ in WT-AL infections (Q2; t = -0.94, p = 0.35), and relative to *per1/2*-AL infections, *per1/2*-RF infections did not differ (Q3; t = 0.66, p = 0.51). The second most supported model also included a treatment × day PI interaction term (model weight 0.17) which we investigate further by comparing peak and cumulative densities.

**FIGURE 2.**
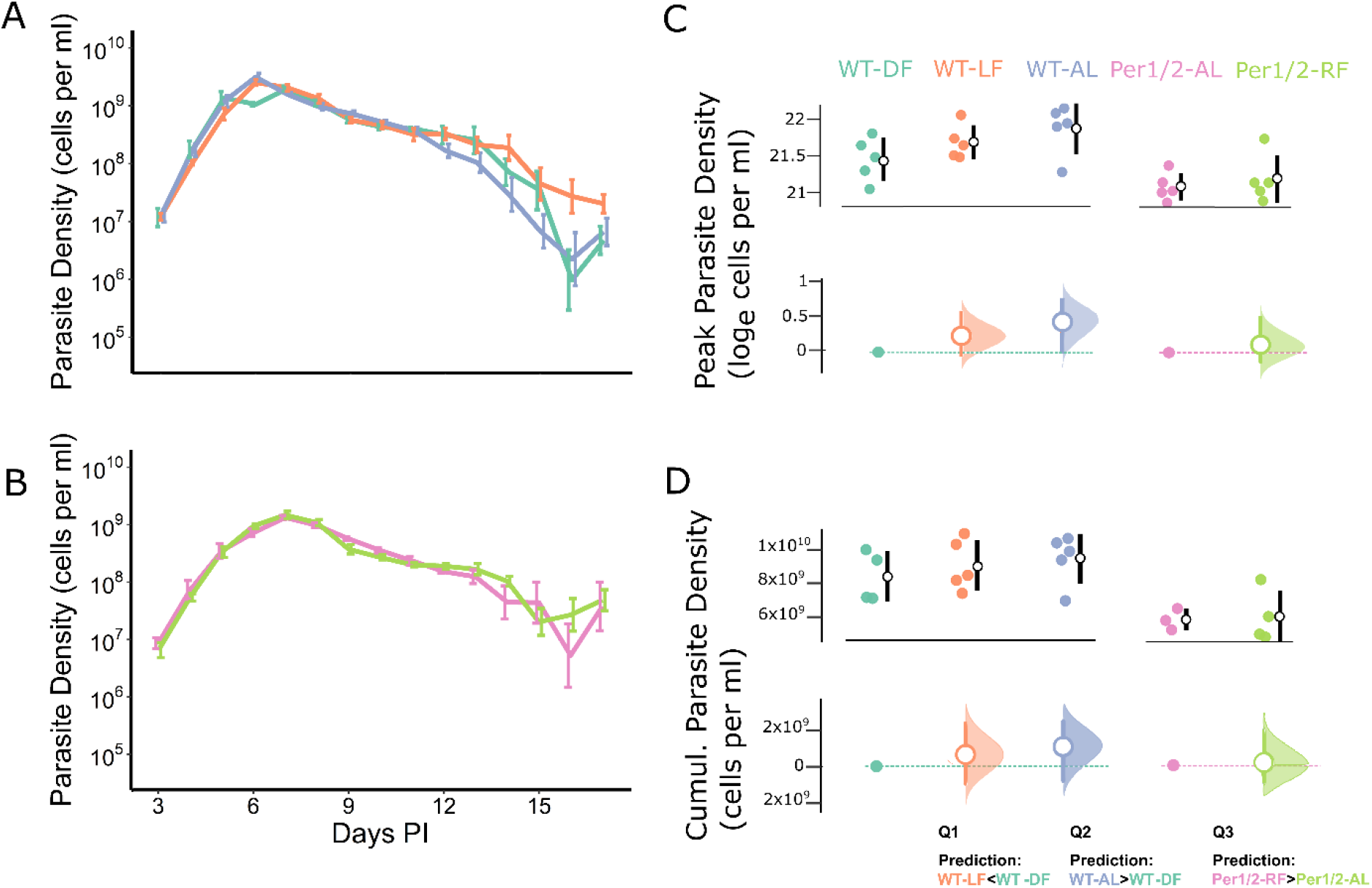
Parasite density metrics. Density dynamics (means ± SE) for (A) infections of wildtype hosts and (B) infections of *per1/2*-null hosts from 3 to 17 days post infection (PI), in which the colours in C correspond to the groups in all figures. Peak parasite densities (C) and cumulative densities (D) by treatment (upper subplot within each panel), along with effect sizes (lower subplot within each panel). Specifically, upper subplots depict (log) peak or cumulative parasite densities per host (coloured points) along with mean ± SE per treatment (white dots and black error bars). Lower subplots depict the effect sizes (mean ± 95% CI difference; white circles and error bars coloured by treatment) of each focal between-treatment comparison (defined by the key questions) of (log) peak or cumulative parasite density. For each comparison, the reference treatment is shown as a point and dotted line (Q1&2, WT-DF, teal; Q3, *per1/2*-AL, pink), and the CIs are derived from nonparametric bootstrap resampling (distribution depicted alongside each error bar). The prediction for each key question is shown below panel D.

The mean (± SE) peak parasite density across treatments was 2.24 × 10^9^ (± 1.83 × 10^8^) cells per ml of blood. The most parsimonious model explaining peak density included a treatment term (model weight = 0.99, delta AICc of null model = 9.4; Table S1). Pairwise comparisons within this model only revealed a difference in the comparison for Q2 (Figure 2C). Specifically, relative to WT-DF infections, the peak did not differ in WT-LF infections (Q1; t = 1.3, p = 0.22), but was 51.4% higher in WT-AL infections (Q2; t = 2.3, p = 0.032; coefficient of log density = 0.41, 95% CI = 0.04-0.79), and relative to *per1/2*-AL infections, the peak was not different in *per1/2*-RF infections (Q3; t = 0.59, p = 0.56).

The mean (± SE) cumulative parasite density across treatments was 7.98 × 10^9^ (± 4.34 × 10^8^) cells per ml of blood. The most parsimonious model explaining cumulative density included a treatment term (model weight = 0.93, delta AICc of null model = 5.1; Table S1). However, only non-focal comparisons (e.g. between WT-DF and *per*1/2-AL) had significant effects, with no significant differences in the pairwise comparisons used to ask Q1, Q2 or Q3 (Figure 2D). Specifically, relative to WT-DF infections, total parasite densities did not differ in WT-LF infections (Q1; t = 0.672, p = 0.511), nor WT-AL infections (Q2; t = 1.112, p = 0.283). Relative to *per1/2*-AL treatment, parasite densities did not differ in the *per*1/2-RF infections (Q3; t = 0.146, p = 0.886).

### 3.2 Gametocyte density

For gametocyte density dynamics, the most parsimonious model included treatment, day PI, and a treatment × day PI interaction (model weight > 0.99, delta AICc of all other models > 34; see Table S2). Gametocyte density dynamics followed similar qualitative patterns across the treatment groups (Figure 3 A, B), with WT-LF infections sustaining higher densities over the first wave (pre 10 days PI), but generally lower in the second wave (post 10 days PI), relative to parasites in WT-DF hosts (Q1, see Table S3), while WT-AL infections were more similar to WT-DF infections throughout (Q2).

**FIGURE 3.**
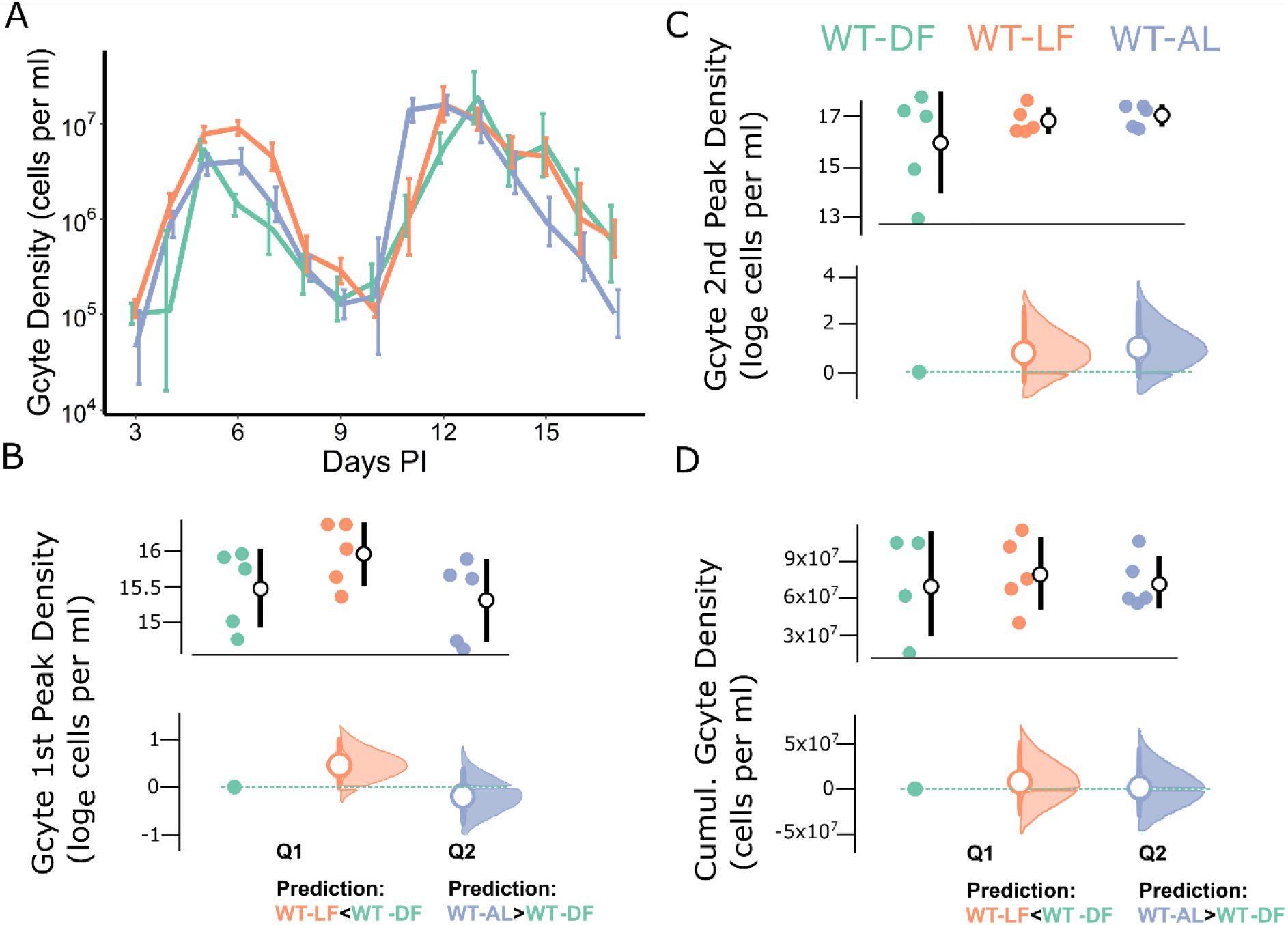
Gametocyte density metrics. Density dynamics (means ± SE) for (A) infections of wildtype hosts from 3 to 17 days post infection (PI), in which the colours in C correspond to the groups in all figures. Peak gametocyte densities of the first (B) and second (C) waves, and cumulative densities (D) by treatment (upper subplot within each panel), along with effect sizes (lower subplot within each panel). Specifically, upper subplots depict (log) peak or cumulative gametocyte densities per host (coloured points) along with mean ± SE per treatment (white dots and black error bars). Lower subplots depict the effect sizes (mean ± 95% CI difference; white circles and error bars coloured by treatment) of each focal between-treatment comparison (defined by the key questions) of (log) peak or cumulative parasite density. For each comparison, the reference treatment, WT-DF, is shown as a teal point and dotted line, and the CIs are derived from nonparametric bootstrap resampling (distribution depicted alongside each error bar). The prediction for each key question is shown below panels B & D.

The mean (± SE) peak gametocyte density was 5.8 × 10^6^ (± 8.0 × 10^5^) cells per ml blood for the first wave and 2.4 × 10^7^ (± 3.6 × 10^6^) cells per ml blood for the second wave. For peak gametocyte density of the first wave, the most parsimonious model included treatment, but only marginally so (model weight = 0.52, delta AICc of null model = 0.18; Figure 3C). However, only non-focal comparisons had significant effects, specifically, relative to WT-DF infections, the early peak was not higher in either WT-LF infections (Q1; t = 0.96, p = 0.35) or WT-AL infections (Q2; t = 0.35, p = 0.73). The most parsimonious model for the peak of the second gametocyte wave did not include treatment (null model weight = 0.98, delta AICc of model with treatment = 7.45).

The mean total (cumulative) gametocyte density across treatments was 6.78 × 10^7^ (± 7.16 × 10^6^) cells per ml blood. The most parsimonious model explaining cumulative density did not include treatment (null model weight = 0.98, delta AICc of model with treatment = 8.38; Figure 3D).

### 3.3 Virulence to hosts

For weight loss, the most parsimonious model included only treatment (model weight = 1.00, delta AICc of null model = 10.96; Figure 4A, B). Pairwise comparisons revealed differences only between the groups used to ask Q2 (Figure 4C). Specifically, compared to WT-DF mice, weight loss did not differ in WT-LF mice (Q1; t = 0.074, p = 0.94), but WT-AL mice lost 35.5% less weight (Q2; t = 2.4, p = 0.026, coefficient = -1.3, 95% CI = -1.84--0.76), and compared to *per1/2*-AL hosts, *per1/2*-RF mice did not differ (Q3; t = 1.0, p = 0.32).

**FIGURE 4.**
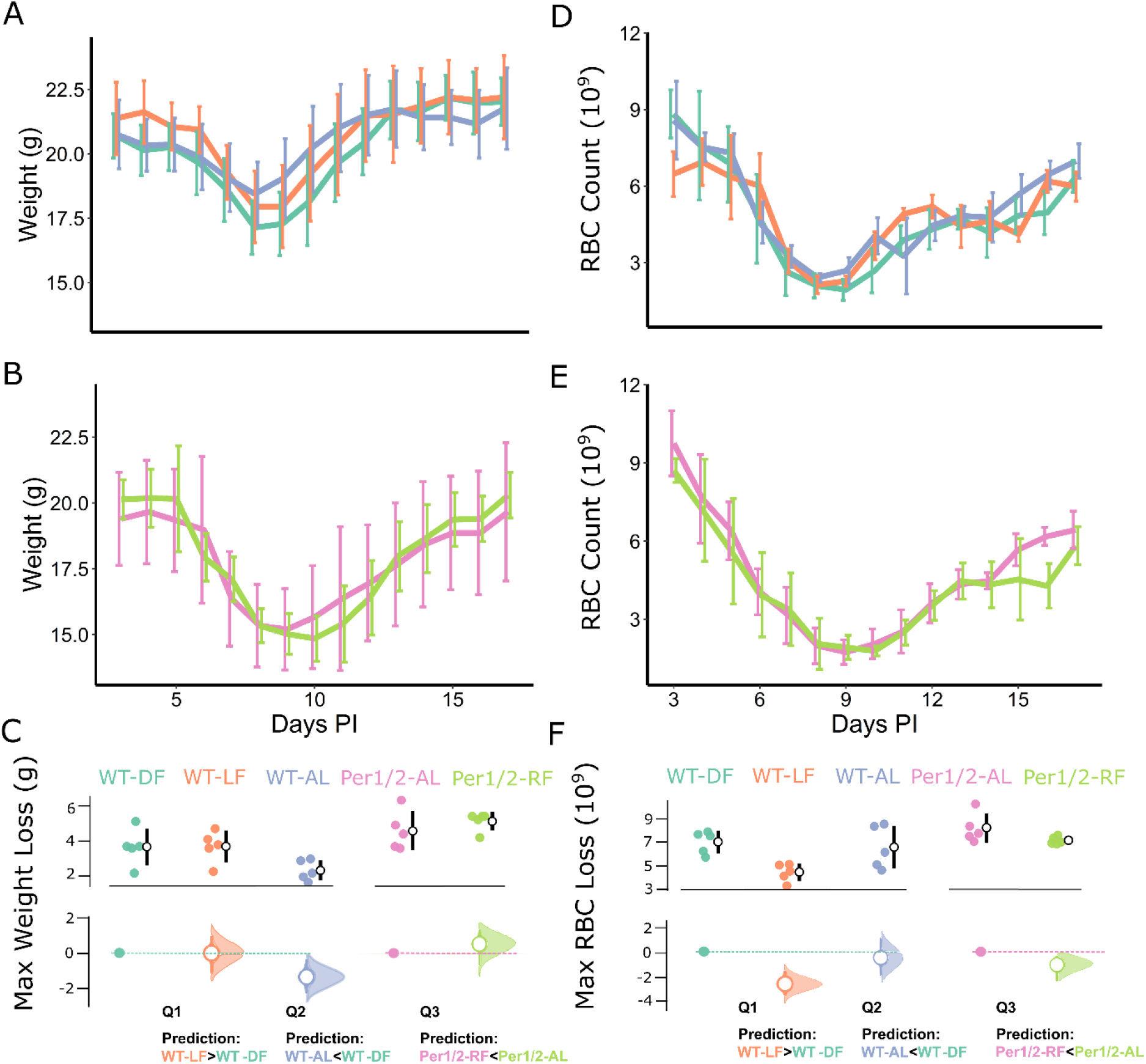
Parasite virulence metrics. Host weight dynamics (means ± SE) for (A) infections of wildtype hosts and (B) infections of *per1/2*-null hosts from 3 to 17 days post infection (PI), in which the colours in C correspond to the groups in all figures. Maximum weight loss per mouse (weight at Day 3 PI minus lowest weight) is shown by treatment (C upper subplot), along with effect sizes (C lower subplot). Host red blood cell (RBC) count dynamics (± SE) for (D) infections of wildtype hosts and (E) infections of *per1/2*-null hosts from 3 to 17 days PI. Maximum RBC loss (in 10^9^ cells per ml) per mouse (RBC count at Day 3 PI minus lowest RBC count) is shown by treatment (F upper subplot), along with effect sizes (F lower subplot). Specifically, upper subplots in C and F depict weight or RBC loss respectively per host (coloured points) along with mean ± SE per treatment (white dots and black error bars). Lower subplots depict the effect sizes (mean ± 95% CI difference; white circles and error bars coloured by treatment)) of each focal between-treatment comparison (defined by the key questions) of weight or RBC loss respectively. For each comparison, the relevant reference treatment is shown as a point and dotted line (Q1&2, WT-DF, teal; Q3, per1/2-AL, pink), and the CIs are derived from nonparametric bootstrap resampling (distribution depicted alongside each error bar). The prediction for each key question is shown below panels C & F.

Finally, for RBC loss, the most parsimonious model did not include treatment (null model weight = 0.93, delta AICc of model with treatment = 5.31; Figure 4D, E, F).

## 4. DISCUSSION

We investigated which host rhythms are the ultimate drivers of the rhythmic replication of malaria parasites, by assessing the impacts of different combinations of host rhythms on within-host survival and transmission potential. In addition, we also tested how perturbations to host rhythms affected the severity of infections. Few focal group comparisons revealed significant differences even though we detected differences between treatments groups for most metrics (6 of 9 model sets), suggesting the analyses had sufficient power to answer our questions. Overall, we reveal more treatment group differences in total parasite than gametocyte density, and that body weight is more sensitive to host rhythm perturbations than anaemia. Specifically, total parasite dynamics followed the same qualitative patterns across treatments, but parasites in WT-LF hosts were able to maintain higher post peak densities than those in WT-DF hosts (Q1) and WT-AL parasites achieved a higher peak than those in WT-DF hosts (Q2). Likewise, gametocyte density dynamics followed qualitatively similar patterns across treatment groups, but with trends for parasites in WT-DF hosts to have a lower first peak than WT-LF infections (Q1) and later second peak than WT-AL infections (Q2), and gametocyte density dropped faster after the second peak in WT-DF than WT-AL infections (Q2). Finally, while red blood cell loss did not differ between the treatment groups, hosts that had the most access to food (WT-AL) lost the least weight (Q2), but internal desynchrony (of light- and food-driven rhythms) did not exacerbate virulence (Q1). Taken together, our results imply that: (Q1) the ultimate driver(s) of parasite rhythms are unlikely to be based on within-host processes driven directly by the light-dark cycle, since parasites (whose IDC rhythm follows host feeding-fasting) did not perform better in any fitness metric when matching the host’s light-driven rhythms; (Q2) parasites benefit when the host feeds in a spread-out-but-rhythmic pattern, since peak parasite density was higher in hosts with a more widely distributed feeding window than hosts with extrinsically-imposed feeding restricted to the night time; and (Q3) whilst imposing rhythmic feeding on clock disrupted hosts is sufficient to generate the IDC rhythm (O’Donnell et al., 2020), this has no apparent ultimate benefit in the absence of the host’s TTFL clock machinery, since parasites did not differ in any metric between clock-disrupted hosts with or without rhythmic feeding.

If light-entrained rhythms are an important ultimate driver of the IDC rhythm, we predicted that parasites would suffer when the IDC is misaligned to light-entrained rhythms (WT-LF). This was not the case, and WT-LF parasites may even perform slightly better at some points during infections. The outputs of light-entrained host clocks usually include the sleep-wake cycle, which is involved in the deployment of T-cells (Besedovsky, Lange, & Born, 2012), some components of innate and adaptive immunity (Carvalho Cabral, Tekade, Stegeman, Olivier, & Cermakian, 2022), and some metabolic and homeostatic processes (e.g. Huang, Ramsey, Marcheva, & Bass, 2011), suggesting these processes have little impact on parasite fitness. Instead, the most parsimonious (but non-mutually exclusive) conclusions of the result of Q1 are that parasites follow host feeding-fasting rhythms to derive benefits from aligning their development with: (i) rhythmicity in blood nutrients derived from food digestion (Skene et al., 2018), (ii) rhythms that more closely follow the phase of host feeding than light cues, including blood oxygen tension (Zhang et al., 2021), body temperature (in small mammals; Abrams & Hammel, 1964), and some immune rhythms (Chen et al., 2022); or (iii) vector activity rhythms, which correlate with nocturnal host feeding-fasting. Aligning with vector rhythms benefits diverse *Plasmodium* species (Pigeault et al., 2018; Schneider et al., 2018), but the fitness consequences of other within-host physiological rhythms remain untested. Alternatively, there may be (iv) no benefits of aligning with environmental rhythms and parasites follow host feeding-fasting simply to ensure an optimal level of synchrony in the IDC. Previous studies suggest that innate immune rhythms do not impose the IDC rhythm by preferentially killing misaligned parasites, though immune cell metabolism may exacerbate the need for the IDC to be aligned with blood nutrient rhythms (Cabral et al., 2022; Hirako et al., 2018; Hunter et al., 2022; Prior et al., 2020). Theory predicts that the optimal level of IDC synchrony is a trade-off between the benefits of being synchronous enough to exploit time-dependent nutrients from the host’s food and the costs of extreme synchrony causing inadvertent competition between parasite cells that are close kin (Greischar et al., 2014; Owolabi, 2023). It is unlikely that WT-LF hosts were less effective at controlling parasites because their health metrics did not differ to WT-DF hosts whose rhythms were not disrupted. Further teasing apart scenarios i-iv and ascertaining their relative importance is empirically very challenging. Confronting parasites with different kinds of rhythms is possible thanks to the tools available for lab mouse models (including conditional clock disruptions). However, experiments must avoid confounding experimental treatments with the costs incurred by parasites altering the IDC rhythm if their alignment to feeding-fasting rhythms is also perturbed.

Given the reliance of parasites on host resources and the potential for high synchrony to be costly, we predicted that parasites derive greater fitness benefits when hosts have access to food throughout the day, compared to hosts whose feeding was restricted to the 12 hour dark phase (albeit with a similar peak feeding time (Q2)). This prediction was supported by parasites in WT-AL hosts achieving a 50% higher peak density. TRF does not typically reduce the amount of food that mice consume per day, even when the window of food availability is much shorter than in our experiment (e.g. 3h; Froy et al., 2006), suggesting that the duration of the feeding window is the key difference between these groups. Furthermore, TRF has broad metabolic impacts, including altering nutrient absorption and temporal expression patterns of various metabolic genes compared to *ad libitum* fed mice (Gallop, Tobin, & Chaix, 2023), suggesting parasites experience significantly better conditions in WT-AL hosts. Spreading out foraging benefited hosts too, ameliorating weight loss despite higher parasite densities. In contrast, WT-DF and WT-AL hosts experience the same degree of anaemia which is surprising since weight loss and anaemia are usually correlated (Timms, Colegrave, Chan, & Read, 2001). This suggests that WT-DF hosts experience a specific difficulty in dealing with malaria-induced appetite reduction when food availability is already limited. This is intriguing, since rodent feeding rhythms plasticly respond to environmental and seasonal changes (Caravaggi et al., 2018; Cohen, Smale, & Kronfeld-Schor, 2009; Tachinardi, Tøien, Valentinuzzi, Buck, & Oda, 2015) and wild house mice have very varied diets (Singleton & Krebs, 2007). Thus, whether WT-DF or WT-AL feeding rhythms – and the costs of infection - better reflect those of wild rodent hosts is likely to depend on the relative consumption of stored food (Singleton & Krebs, 2007) and rhythmically available forage (e.g. due to ambient temperatures and energetic efficiency; Hut, Pilorz, Boerema, Strijkstra, & Daan, 2011) in their evolutionary history.

Our third question (Q3) specifically considered whether feeding-fasting rhythms act as an ultimate driver (as well as a proximate cue) for the IDC rhythm (Prior et al., 2021). Directly testing this hypothesis requires misaligning parasites to host feeding-fasting rhythms and quantifying the fitness consequences. Unfortunately, it is not possible to prevent parasites from rescheduling the IDC to realign to feeding-fasting rhythms (which can occur within 5 cycles, O’Donnell et al., 2022; O’Donnell et al., 2020; O’Donnell et al., 2011). Thus, our finding that parasite performance did not differ between infections in clock-disrupted hosts, regardless of whether a feeding rhythm was imposed via TRF (i.e. *per*1/2-AL vs *per*1/2-RF) has several possible (non-mutually exclusive) interpretations. First, host feeding-fasting rhythms are a proximate but not ultimate driver of the IDC rhythm, allowing parasites to align to other (typically correlated) rhythms or to achieve the optimal level of synchrony (discussed in Q1, above). However, we propose that the intrinsic benefits hypothesis is least likely to explain the IDC rhythm because it predicts that synchrony is adaptive regardless of resource availability (i.e. *per*1/2-RF parasites should have outperformed *per*1/2-AL parasites, which they did not). Second, there are benefits of aligning specifically with feeding-fasting rhythms but these are offset by other costs. For example, parasites in *per*1/2-RF hosts may benefit from aligning IDC stages with the availability of the nutrients they need but the TRF window might have increased synchrony to a costly level (analogous to why parasites in WT-AL hosts may perform better than those in WT-DF hosts).

Alternatively, parasites in *per*1/2-AL hosts may not experience nutrient limitation but may suffer from a loss of intrinsic benefits (if synchrony alone is adaptive). These different costs and benefits may coincidently result in no net differences between treatment groups. Third, the ultimate driver of the IDC rhythm may be a host rhythm that requires input from both feeding-fasting rhythms and the TTFL clock, which was not experienced by parasites in *per*1/2-RF hosts. This scenario suggests that only a TTFL-mediated component of feeding-fasting rhythms acts as an ultimate driver for IDC rhythms and that time cue(s) such as isoleucine (which can be rhythmic in the absence of the TTFL) function as a proxy. Mechanistically, this fits with recent models of peripheral rhythms in mammals, whereby feeding drives some downstream physiological outputs directly, but others are mediated by TTFL clocks in the liver and other peripheral organs, as well as the central pacemaker in the SCN (Zhang et al., 2020). For example, both rhythmic feeding and a functioning TTFL clock are needed to generate rhythmic gene expression of some metabolic genes involved in lipogenesis and glycogenesis in the liver (Greenwell et al., 2019; Vollmers et al., 2009), and so these or similar metabolites may be candidate ultimate drivers of parasite rhythmicity.

## Conclusions

Our study complements the increasing body of work focussing on ‘how’ *Plasmodium* parasites set the schedule of the IDC rhythm by asking ‘why’ this rhythm is adaptive. Identifying selective drivers is a challenge for a trait that does not have the benefit of, for example, the well-established theoretical literature on adaptation that the field of life history evolution benefits from. Nonetheless, our results eliminate purely light-driven rhythmic host processes as being the selective, ultimate, driver of parasite rhythmicity. Further work is required to explore the new hypothesis we propose; that feeding-fasting rhythms are a selective driver but only when mediated by TTFL clocks. Testing this, along with the ultimate roles of other rhythms, such as oxygen tension and vector activity, would be facilitated by knowing the molecular mechanism(s) that underpin the timing and synchrony of the IDC. For example, blocking parasites’ ability to sense time or alter the IDC schedule would stabilise misalignment to host feeding-fasting rhythms, enabling fitness consequences to be directly assessed. While our experimental design improves on previous studies of fitness consequences by considering whole infections rather than a few IDCs, the adaptive value of rhythms in other taxa have been most clearly demonstrated in stressful conditions such as competition (Dodd et al., 2005; Fleury, Allemand, Vavre, Fouillet, & Bouletreau, 2000; Ouyang, Andersson, Kondo, Golden, & Johnson, 1998). Within-host competition is frequently experienced by *Plasmodium* species and experimentally tractable, as is manipulating overall resource availability via the diet of hosts. Overall, the relatively modest impacts of host rhythm manipulations on within-host parasite fitness metrics emphasises the role of rhythmic transmission opportunities as a putative ultimate driver, as proposed to explain rhythms in other parasite taxa (e.g. *Wucheria*, Hawking, 1967; *Schistosoma*, Mouahid et al., 2012; and *Isospora*, Martinaud, Billaudelle, & Moreau, 2009). Finally, if human malaria parasites use proximate time cues to align with other rhythms (that ultimately select for the IDC rhythm), this opens up the potential to develop interventions that act as ecological traps by coercing parasites into adopting a sub optimal IDC schedule that reduces transmission and dampens virulence.

## STATEMENTS

The authors declare no conflicts of interest. All procedures were carried out in accordance with the UK Home Office regulations (Animals Scientific Procedures Act 1986; SI 2012/3039) and approved by the ethical review panel at the University of Edinburgh. Data for this study are available at [*to be completed after manuscript is accepted for publication*].

## ACKNOWLEDGEMENTS

We thank Ronnie Mooney, Aliz Owolabi, Petra Schneider, and Mary Westwood for assistance, and Giles K P Barra for technical advice. Funding was provided by Wellcome (202769/Z/16/Z) and the Royal Society (URF\R\180020).

## supplementary material

**Table S1.** Parasite density models. The three table sections show model selection for, respectively, Log(parasite density), Log(parasite peak density) or Cumulative parasite density as response variables, with Treatment as a fixed factor. Day PI (Day, as a fixed factor), Day × Treatment interaction, and Mouse ID (as random intercepts) were also included in the full parasite density model. Degrees of freedom (Df), log likelihood (Log lik), ΔAICc (the most parsimonious model is indicated with 0 in bold), and model weight (weight) are shown for each analysis. The coefficient (coef) and standard error of the mean (SE) for the three key questions are given for the most parsimonious model, with significant differences in bold. Specifically, the coefficients are as follows: Q1 = WT-LF treatment, using WT-DF as reference level; Q2 = WT-AL treatment, using WT-DF as reference level; Q3 = Per1/2-RF treatment, using Per1/2-AL as reference level.

| Model | Df | Log lik | $\Delta AICc$ | weight | Q1 coef<br>±SE | Q2 coef<br>±SE | Q3 coef<br>±SE |
| --- | --- | --- | --- | --- | --- | --- | --- |
| **Log(Density) ~ Day + Treatment + (1 \| MouseID)** | **21** | **-441.60** | **0** | **0.796** | **0.36 ± 0.15** | -0.14 ± 0.15 | 0.11 ± 0.16 |
| Log(Density) ~ Day + Treatment + Day x Treat + (1 \| MouseID) | 77 | -364.75 | 3.08 | 0.170 |  |  |  |
| Log(Density) ~ Day + (1 \| MouseID) | 17 | -449.27 | 6.36 | 0.033 |  |  |  |
| Log(Density) ~ Treatment + (1 \| MouseID) | 7 | -482.82 | 51.90 | 0 |  |  |  |
| Log(Density) ~ 1 + (1 \| MouseID) (null) | 3 | -490.13 | 58.24 | 0 |  |  |  |
| **Log(Peak Height) ~ Treatment** | **6** | **-1.27** | **0** | **0.991** | 0.23 ± 0.18 | **0.41 ± 0.18** | 0.11 ± 0.18 |
| Log(Peak Height) ~ 1 (null) | 2 | -12.01 | 9.36 | 0.009 |  |  |  |
| **Cumul. Density ~ Treatment** | **6** | **-496.73** | **0** | **0.927** | (6.5 ± 9.6) x 10<sup>8</sup> | (10.7 ± 9.6) x 10<sup>8</sup> | (-1.6 ± 11.0) x 10<sup>8</sup> |
| Cumul. Density ~ 1 (null) | 2 | -478.94 | 5.07 | 0.073 |  |  |  |

**Table S2.**
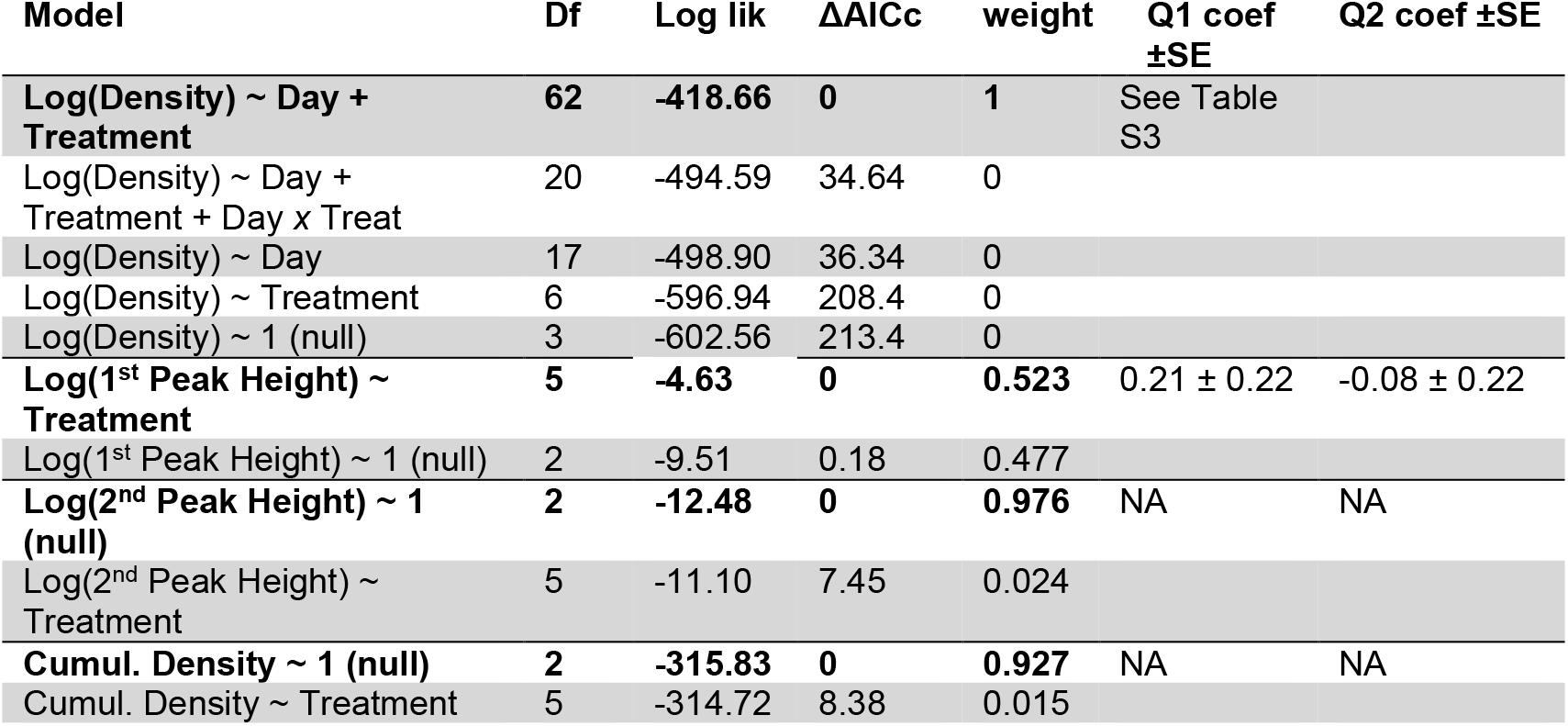
Gametocyte density models. The four table sections show model selection for, Log(gametocyte density), Log(gametocyte peak density for the 1^st^ wave), Log(gametocyte peak density for the 2^nd^ wave), or Cumulative gametocyte density as response variables, with Treatment as a fixed factor. Day PI (Day, as a fixed factor), Day × Treatment interaction, and Mouse ID (as random intercepts) were also included in the full gametocyte density model. Degrees of freedom (Df), log likelihood (Log lik), ΔAICc (the most parsimonious model is indicated with 0 in bold), and model weight (weight) are shown for each analysis.. For the peak and cumulative density models, the coefficient (coef) and standard error of the mean (SE) for two of the key questions are given for the most parsimonious model. Specifically, the coefficients given are as follows: Q1 = WT-LF treatment, using WT-DF as reference level; Q2 = WT-AL treatment, using WT-DF as reference level.

| Model | Df | Log lik | ΔAICc | weight | Q1 coef<br>±SE | Q2 coef ±SE |
| --- | --- | --- | --- | --- | --- | --- |
| **Log(Density) ~ Day + Treatment** | **62** | **-418.66** | **0** | **1** | See Table S3 |  |
| Log(Density) ~ Day + Treatment + Day x Treat | 20 | -494.59 | 34.64 | 0 |  |  |
| Log(Density) ~ Day | 17 | -498.90 | 36.34 | 0 |  |  |
| Log(Density) ~ Treatment | 6 | -596.94 | 208.4 | 0 |  |  |
| Log(Density) ~ 1 (null) | 3 | -602.56 | 213.4 | 0 |  |  |
| **Log(1<sup>st</sup> Peak Height) ~ Treatment** | **5** | **-4.63** | **0** | **0.523** | 0.21 ± 0.22 | -0.08 ± 0.22 |
| Log(1<sup>st</sup> Peak Height) ~ 1 (null) | 2 | -9.51 | 0.18 | 0.477 |  |  |
| **Log(2<sup>nd</sup> Peak Height) ~ 1 (null)** | **2** | **-12.48** | **0** | **0.976** | NA | NA |
| Log(2<sup>nd</sup> Peak Height) ~ Treatment | 5 | -11.10 | 7.45 | 0.024 |  |  |
| **Cumul. Density ~ 1 (null)** | **2** | **-315.83** | **0** | **0.927** | NA | NA |
| Cumul. Density ~ Treatment | 5 | -314.72 | 8.38 | 0.015 |  |  |

**Table S3.** Gametocyte density dynamics from the most parsimonious model of parasite density, including a Day × Treatment interaction term. Coefficients and standard errors corresponding to two of the key questions; i.e. the WT-LF (Q1) and WT-AL (Q2) groups, relative to the reference group WT-DF. The intercept (Day 3 for WT-DF) is given.

| Day | Reference coef ±SE<br>(WT-DF) | Q1 coef ±SE | Q2 coef ±SE |
| --- | --- | --- | --- |
| **3** | 11.54 ± 0.61 (intercept) | 0.12 ± 0.87 | -0.82 ± 0.87 |
| **4** | 0.08 ± 0.83 | 2.41 ± 1.17 | 2.90 ± 1.17 |
| **5** | 3.96 ± 0.83 | 0.25 ± 1.17 | 0.47 ± 1.17 |
| **6** | 2.63 ± 0.83 | 1.73 ± 1.17 | 1.87 ± 1.17 |
| **7** | 2.03 ± 0.83 | 1.63 ± 1.17 | 1.42 ± 1.17 |
| **8** | 0.95 ± 0.83 | 0.37 ± 1.17 | 0.95 ± 1.17 |
| **9** | 0.36 ± 0.83 | 0.55 ± 1.17 | 0.69 ± 1.17 |
| **10** | 0.76 ± 0.83 | -0.81 ± 1.17 | 0.48 ± 1.17 |
| **11** | 2.41 ± 0.88 | -0.19 ± 1.21 | 3.33 ± 1.21 |
| **12** | 4.05 ± 0.88 | 0.89 ± 1.21 | 1.81 ± 1.21 |
| **13** | 5.27 ± 0.88 | -0.70 ± 1.21 | 0.18 ± 1.21 |
| **14** | 3.74 ± 0.88 | 0.03 ± 1.21 | 0.48 ± 1.21 |
| **15** | 4.11 ± 0.88 | -0.44 ± 1.21 | -1.07 ± 1.21 |
| **16** | 2.75 ± 0.88 | -0.58 ± 1.21 | -0.56 ± 1.21 |
| **17** | 1.73 ± 0.88 | -0.04 ± 1.21 | -0.91 ± 1.21 |

**Table S4.** Host virulence models. The two table sections show model selection for, Weight Loss or RBC loss as response variables, with Treatment as a fixed factorDegrees of freedom (Df), log likelihood (Log lik), ΔAICc (with most parsimonious model shown with 0 in bold), and model weight (weight) are shown. The coefficient (coef) and standard error of the mean (SE) for two of the key questions are given for the most parsimonious model, with significant differences in bold. Specifically, the coefficients given are as follows: Q1 = WT-LF treatment, using WT-DF as reference level; Q2 = WT-AL treatment, using WT-DF as reference level.

| Model | Df | Log lik | $\Delta AICc$ | weight | Q1 coef<br>±SE | Q2 coef<br>±SE | Q3 coef<br>±SE |
| --- | --- | --- | --- | --- | --- | --- | --- |
| **Weight loss ~ Treatment** | **6** | **-28.72** | **0** | **0.996** | 0.04 ± 0.54 | -1.3 ± 0.54 | 0.54 ± 0.54 |
| Weight loss ~ 1 (null) | 2 | -40.26 | 10.96 | 0.004 |  |  |  |
| **RBC loss ~ 1 (null)** | **2** | **-14.27** | **0** | **0.934** | NA | NA | NA |
| RBC loss ~ Treatment | 6 | -10.86 | 5.31 | 0.066 |  |  |  |

